# Molecular features of exceptional response to neoadjuvant anti-androgen therapy in high-risk localized prostate cancer

**DOI:** 10.1101/2021.04.20.440657

**Authors:** Alok K. Tewari, Alexander T.M. Cheung, Jett Crowdis, Jake R. Conway, Sabrina Y. Camp, Stephanie A. Wankowicz, Dimitri G. Livitz, Jihye Park, Rosina T. Lis, Alice Boosma-Moody, Meng Xiao He, Saud H. AlDubayan, Zhenwei Zhang, Rana R. McKay, Ignaty Leshchiner, Myles Brown, Steve Balk, Gad Getz, Mary-Ellen Taplin, Eliezer M. Van Allen

**Author notes:** These authors contributed equally (Alok K. Tewari, Alexander T.M. Cheung). These authors contributed equally (Mary-Ellen Taplin, Eliezer M. Van Allen).

## Abstract

High-risk localized prostate cancer (HRLPC) is associated with a substantial risk of recurrence and prostate cancer-specific mortality^1^. Recent clinical trials have shown that intensifying anti-androgen therapies administered prior to prostatectomy can induce pathologic complete responses (pCR) or minimal residual disease (MRD) (<5 mm), together termed exceptional response, although the molecular determinants of these clinical outcomes are largely unknown. Here, we performed whole exome (WES) and whole transcriptome sequencing (RNA-seq) on pre-treatment multi-regional tumor biopsies from exceptional responders (ER: pCR and MRD patients) and non-responders (NR: pathologic T3 or lymph node positive disease) treated with intensive anti-androgen therapies prior to prostatectomy. *SPOP* mutation and *SPOPL* copy number loss were exclusively observed in ER, while *TP53* mutation and *PTEN* copy number loss were exclusively observed in NR. These alterations were clonal in all tumor phylogenies per patient. Additionally, transcriptional programs involving androgen signaling and TGFβ signaling were enriched in ER and NR, respectively. The presence of these alterations in routine biopsies from patients with HRLPC may inform the prospective identification of responders to neoadjuvant anti-androgen therapies to improve clinical outcomes and stratify other patients to alternative biologically informed treatment strategies.

## Introduction

Over 90% of the 191,930 estimated cases of prostate cancer (PCa) in 2020 will present as localized disease^2^, with approximately 15% of these cases defined as high-risk for recurrence^2–4^. Though HRLPC is often curable by surgery alone, or with combined radiation plus androgen-deprivation therapy (ADT), the risk of progressive disease and long-term prostate cancer specific mortality approaches 40%^1,5^. In other malignancies^6–11^, neoadjuvant systemic therapy results in pCR or MRD in a subset of patients and is associated with improved overall survival. Neoadjuvant treatment strategies centered on androgen signaling pathways are being tested in HRLPC, since PCa is generally dependent on androgen and its cellular receptor, the androgen receptor (*AR*).

Neoadjuvant trials of androgen-pathway inhibitors (APIs) performed in the 1990s were limited by heterogeneous patient-risk groups predominated by low risk PCa, inadequate antagonism of the *AR* and reductions in intra-prostatic androgen signaling, and lack of long-term survival data^12–17^. Thus, contemporary neoadjuvant efforts for treating HRLPC have focused on using more intensive treatment strategies that combine conventional ADT with newer APIs such as the androgen-synthesis inhibitor abiraterone^18–21^, and the *AR* antagonists enzalutamide^22–24^ or apalutamide^25,26^. Our recent trial of intensive neoadjuvant treatment with enzalutamide, ADT, abiraterone, and prednisone (ELAP) for 6 months prior to prostatectomy induced pCR or MRD in 30% of patients^27^ and pCR/MRD is associated with freedom from biochemical recurrence (M. Taplin, communication, and^28^). Though studies have identified genomic associations with response to API in advanced PCa^29,30^ as well as prognostic molecular features of aggressive localized disease^31,32^, the underlying biology and molecular mediators of response to intensive neoadjuvant API remains largely unknown. Given that integrative molecular features have been implicated in selective ADT response in the advanced prostate cancer setting, we hypothesized that genomic and transcriptional properties may be operant and coordinated in HRLPC treated with intensive neoadjuvant therapy

## Results

### Whole exome analysis of ER and NR patient samples

We performed multi-regional WES and RNA-seq profiling of pre-treatment biopsies from 32 patients enrolled on neoadjuvant trials of intensive API with either ELAP or ADT with apalutamide and abiraterone (AAP) (Figure 1A). A total of 46 WES samples from 24 patients (21 high-risk and 3 unfavorable intermediate risk by NCCN criteria) passed quality control metrics and were included in the final cohort (Figure 1A, Figure S1A, Table S1, Methods). There were no significant differences in patient age, prostate specific antigen (PSA) or histopathology (Gleason score) between the 13 ER and 11 NR patients (Wilcoxon rank sum, Table 1). Of the 46 samples, 43 had a matched RNA-seq sample extracted from the same tissue that passed quality control (Methods). The median WES coverage for tumor and germline samples was 158x and 85x, respectively. Tumor purity varied across the samples from 0.21 to 0.84 but did not differ between response groups (*P* = 0.2793, Wilcoxon rank sum, Table S1). The median purity-corrected tumor ploidy in each response group was 2.1, as one NR tumor (27_T3) had a whole genome doubling event. There was no significant difference in median tumor mutational burden (TMB) between ER and NR groups (*P* = 0.093, Wilcoxon rank sum, Figure 1B, left panel). Similarly, no significant difference in the proportion of genome altered (PGA) was observed between the groups (*P* = 0.502, Kolmogorov-Smirnov, Figure 1B, right panel).

**Figure 1:**
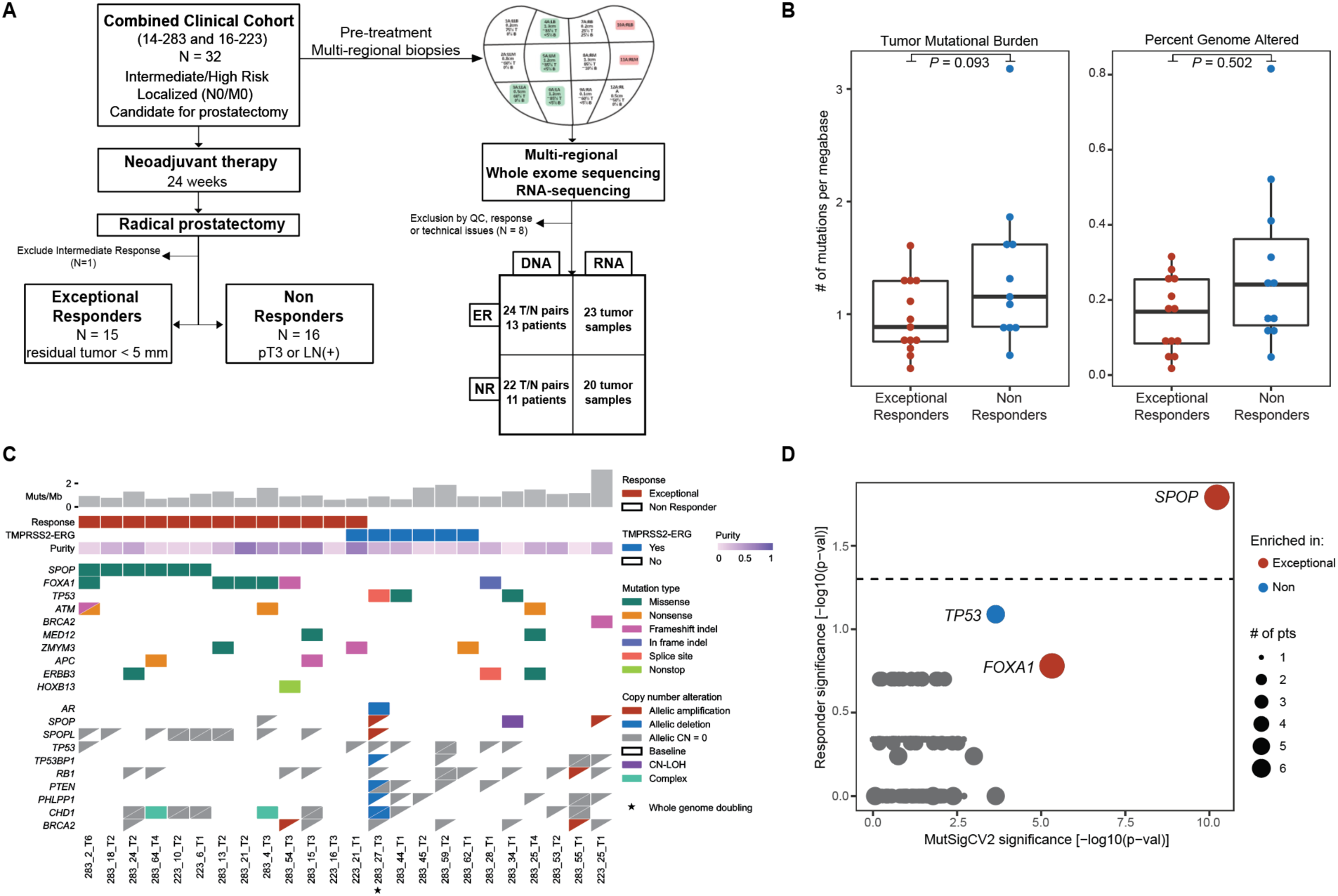
Genomic features of response to neoadjuvant API. (a) Study design integrating WES and RNA-seq pre-treatment biopsies from two clinical trial protocols of ELAP (14-283) or AAP (16-223) (b) Comparison of TMB (*P* = 0.093, Wilcoxon rank-sum) and PGA (*P* = 0.502, Kolmogorov-Smirnov) between ER (n = 13) and NR (n=11) patients. (c) Co-mutation^88^ plot of the highest purity sample per patient illustrating select single nucleotide and insertion/deletion events, as well as CNA and *TMPRSS2-ERG* RNA fusion status. Each row represents the mutation or copy number status for the indicated gene, and each column represents a patient sample. Copy number calls are allelic, with the status of each allele indicated by a triangle. In cases with whole genome doubling (27_T3), it is possible for one allele to be amplified and one or both of the other allele to be lost. *AR* is on the X chromosome with only a single copy in men and is thus represented by a solid box for copy number status. Complex indicates that a segment breakpoint occurred within a gene, creating conflicting copy number. Baseline indicates the default copy number status of a diploid genome with 1 copy of each allele and corresponds to 2 copies per allele in the case of whole genome doubling. Mutations are not annotated by allele. Each gene shown had only one unique mutation per sample, with the exception of *ATM*, which had two. (d) Enrichment in non-synonymous mutations between ER and NR patient groups (y-axis, Fisher’s exact test) in relation to mutational significance across the entire cohort (x-axis). The dashed line indicates a *P*-value threshold of 0.05, and circle size reflects the number of patients which the indicated alteration is present in.

**Table 1:** Clinical cohort characteristics. *P*-values for differences in age, baseline PSA, and Gleason score calculated using Wilcoxon rank-sum test.

|  | Total (N = 24) | Responders (N = 13) | Non Responders (N = 11) |  |
| --- | --- | --- | --- | --- |
| **Age at Diagnosis** | 62.36 ± 6.2 | 62.96 ± 7.5 | 63.73 ± 4.6 | P = 0.862 |
| **Race** |  |  |  |  |
| White | 21 | 10 | 11 |  |
| Asian | 2 | 2 | 0 |  |
| Unknown | 1 | 1 | 0 |  |
| **PSA (ng/mL)** | 5.84 ± 10.5 | 5.92 ± 10.5 | 5.2 ± 11.0 | P = 0.5691 |
| **Clinical Staging, n** |  |  |  |  |
| T1 | 6 | 4 | 2 |  |
| T2 | 13 | 7 | 6 |  |
| T3 | 5 | 2 | 3 |  |
| **Gleason Score** | 8 ± 0.9 | 8 ± 0.9 | 9 ± 0.9 | P = 0.1789 |

Examination of genomic alterations within this cohort (Table S2) revealed *SPOP* missense mutation (6/13 patients), *SPOP* copy number loss (1/13 patients), and *SPOPL* copy number loss (8/13 patients) exclusively in ER. Conversely, *TP53* mutation (3/11 patients), *PTEN* loss (4/11 patients), *PHLPP1* loss (6/11 patients), and *AR* deletion (1/11 patients) were exclusively observed in NR. *TMPRSS2-ERG* fusion transcripts were detected in 6 patients, 5 of whom were NR (Figure 1C, Figure S1B). Copy number alterations in *RB1* and *TP53* were present in both ER and NR. Deletion of chromosome 5q21.1, which contains *CHD1*, was significantly enriched in ER by GISTIC2.0^33^ (*Q* = 0.053, Table S3, Methods). Of all genes with at least one potentially damaging somatic alteration, only *SPOP* and *FOXA1* were significantly mutated across our cohort by MutSigCV2^34^ (*Q* < 0.05, Methods), and only *SPOP* mutation was significantly enriched in ER (*P =* 0.016, Fisher’s exact test, Figure 1D, Table S4, Methods).

We then compared our cohort with an untreated localized PCa molecular cohort (The Cancer Genome Atlas (TCGA) study of localized PCa^35^). We found that our cohort had a significantly lower TMB (*P* = 2×10^-6^, Wilcoxon rank sum) but no difference in PGA (*P* = 0.237, Kolmogorov-Smirnov) relative to the TCGA dataset (Figure S1C). TCGA samples with *SPOP* mutation also had *SPOPL* copy number loss and *APC* mutation (*P* < 0.001, Fisher’s exact test; co-occurrence), and lacked *PTEN* copy number loss or *TMPRSS2-ERG* fusion (*P* = 0.046 and *P* < 0.0001, respectively, Fisher’s exact test; mutual exclusivity; Figure S1D). *SPOP* mutations in our cohort were also present in the TCGA cohort, though we did identify a *TP53* splice site mutation (chr17:7577497A>C) not present in the TCGA cohort (Figure S1E). Thus, ER and NR cohorts were associated with recurrent mutations in *SPOP* and *TP53*, respectively.

### Clonal architecture of ER and NR tumors

WES studies of localized PCa have identified substantial intra-tumoral heterogeneity between tumor foci^36–40^. We therefore evaluated pre-treatment multifocal biopsies to interrogate the association between intra-tumoral heterogeneity and response as well as the clonal architecture of genes associated with response to neoadjuvant API. A total of 14 patients (8 ER, 6 NR) had multiple biopsies available for analysis. A median of 62.5% (range: 18.3-96.6) of non-synonymous mutations were common across at least two biopsies within each of these patients.

To evaluate the evolutionary relationship between intraprostatic tumor foci, we utilized phylogicNDT^41,42^ to cluster mutations by cancer-cell fraction (CCF) and build phylogenetic trees of tumor evolution in patients with multifocal biopsies (Methods). Of the 14 patients with multifocal biopsies available, 12 had tumor foci that shared a truncal mutational cluster. It is possible that the two patients for which the mutational cluster with the highest average CCF was not present in all biopsies (283_21 and 283_54, evidenced by low CCF of the assigned truncal cluster in some of the biopsy sites), the tumor foci sampled evolved independently and were not clonally related. Importantly, amongst the 3 *SPOP*-mutant patients with multifocal biopsies, *SPOP* mutation was a truncal event, present in all samples at high CCF (Figure 2A). We observed a similar finding in the 2 *TP53*-mutant patients with multifocal biopsies (Figure 2B). Conversely, though we observed many truncal mutations in other PCa genes, this was not always the case, such as a *PTEN* mutation in patient 283_21 present at high CCF in only one tumor focus (Figure 2B).

**Figure 2:**
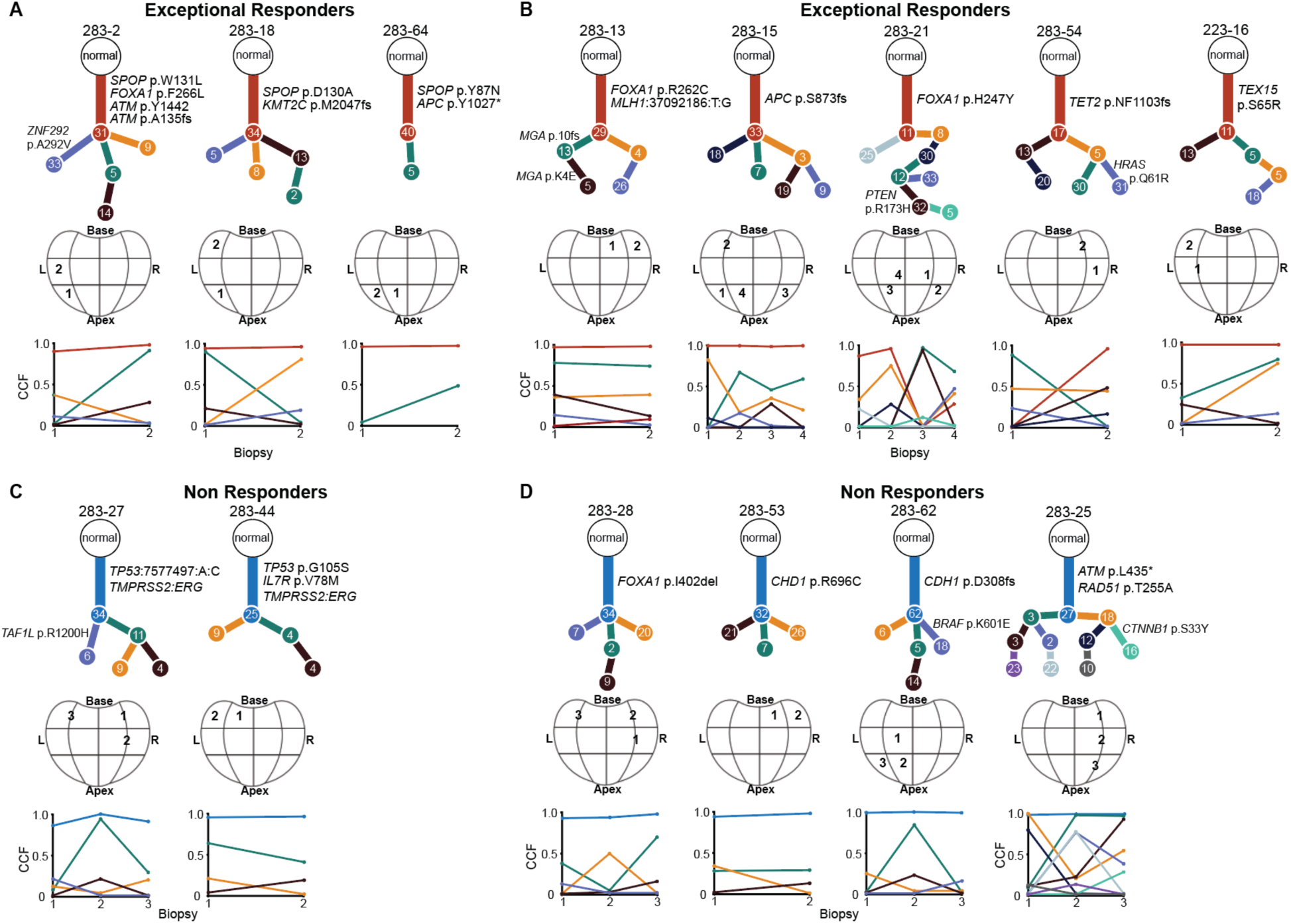
Phylogenetic trees of select ER and NR patients reveal truncal alterations in *SPOP* and *TP53*. **(A)** *PhylogicNDT BuildTree* evolutionary trees of ER patients with *SPOP* mutation. Known prostate cancer associated genes are labelled according to their position in the tree. Each distinct mutational cluster is illustrated with a different cluster, with the number of mutations in each cluster indicated in each node. No information is reflected in the length of each branch. Biopsy location is approximated in the 12-grid prostate representation below each tree, and the average cancer cell fraction of each mutational cluster within each biopsy sample is illustrated below the grid. **(B)** Phylogenetic trees of ER patients lacking *SPOP* mutation generally reveal truncal mutations in known prostate cancer associated genes. **(C)** Phylogenetic trees of two NR patients with *TP53* mutation. *TMPRSS2-ERG* fusion status is labelled as truncal based on its detection in all samples from the indicated patient. **(D)** Phylogenetic trees of NR patients lacking *TP53* mutation, including patient 283-25 with more complex clonal architecture associated with multiple missense mutations in DNA damage repair genes.

### Mutational signatures and clinical response

Since prior reports have suggested *SPOP* mutation may lead to genomic instability via modulation of homology-directed repair of DNA double-stranded breaks^43,44^, we then explored whether response was associated with homologous recombination deficiency (HRD) or other genomic alteration patterns. There was no difference in TMB or PGA between *SPOP-*mutant and non-mutant samples within ER (*P* = 0.76 for TMB, *P =* 0.68 for PGA, Wilcoxon rank sum) or across the cohort (*P* = 0.62 for TMB, *P =* 0.77 for PGA, Wilcoxon rank sum). Furthermore, all samples were microsatellite stable (MSIsensor^45^ scores < 0.5).

To assess for evidence of specific mutational processes and HRD in our cohort, we utilized mutation-based (deconstructSigs^46^ and SigMA^47^) and copy number-based (scarHRD^48^) methods (Methods). Consistent with prior reports^47,49^, clock-like signature 1 was the predominant mutational signature present across all samples. No COSMIC^50^ mutational signature was significantly enriched in either response group (Figure S2A, Methods). We observed higher scarHRD scores in NR, suggestive of more HRD-associated genomic structural alterations relative to ER samples (*P* = 0.098, Wilcoxon rank sum, Figure S2B, Table S5). Two NR had biopsy scarHRD scores ≥ 42, a threshold considered suggestive of HRD in preclinical studies^51,52^ as well as by the Myriad CLIA-certified myChoice HRD score. One of these patients, 283_25, harbored somatic mutations in multiple DNA damage associated genes such as *ATM, RAD51*, *FANCA*, and *FANCL* amongst others (Figure 2D, Table S2) whereas the other patient, 283_55, had no detected alteration in a DNA damage repair gene. SigMA, which is optimized for samples with lower overall mutational burden, detected the presence of the HRD-associated signature 3 in multiple samples in the ER and NR categories, including patient 223_25 who harbored both germline and somatic mutations in *BRCA2*. Beyond the *BRCA2* germline mutation in patient 223_25, no other germline alteration within a curated set of DNA damage repair genes was observed in NR (Fisher’s exact test, Table S6).

### Transcriptome sequencing identifies androgen signaling in ER and TGFβ signaling in NR

In addition to evaluating genomic events mediating selective response, we also assessed transcriptional programs associated with response to neoadjuvant API in 43 pre-treatment RNA-seq samples available from the same tumor foci as the WES samples. Gene set enrichment analysis (GSEA)^53^ of MSigDB hallmark gene sets^54^ identified 7 and 11 gene sets enriched in ER and NR at a false-discovery rate < 0.25, respectively (Figure S3A, Methods). Specifically, the androgen signaling gene set was enriched in ER, and gene sets consistent with inflammatory pathways and TGFβ signaling were enriched in NR (Figure 3A, Figure S3A). We next utilized single-sample GSEA^55^ (ssGSEA) to assess if the observed up-regulated androgen signaling in ER tumors are driven by *SPOP* mutation. As expected, ssGSEA scores for the androgen signaling gene set were significantly higher in ER compared to NR (*P* = 0.003, Wilcoxon rank sum); however, there was no significant difference in androgen signaling enrichment between *SPOP*-mutant or wild-type ER (Figure 3B). We next used multiple approaches (Methods) to identify a consensus list of genes differentially expressed by response (Figure 3C, Figure S3B). We identified 136 genes up-regulated in ER and 21 genes up-regulated in NR. ER up-regulated genes included those regulated by *AR* such as *CAMKK2* and *SLC45A3*, *mTOR*, and *ANPEP*, whose expression loss has been proposed as a prognostic marker in localized PCa^56^ (Figure S3C). Genes up-regulated in non-responders had overall expression levels less than those up-regulated in ER tumors, and include *PLA2G7*, a gene associated with PCa cell migration and invasion^57^ (Figure S3D).

**Figure 3:**
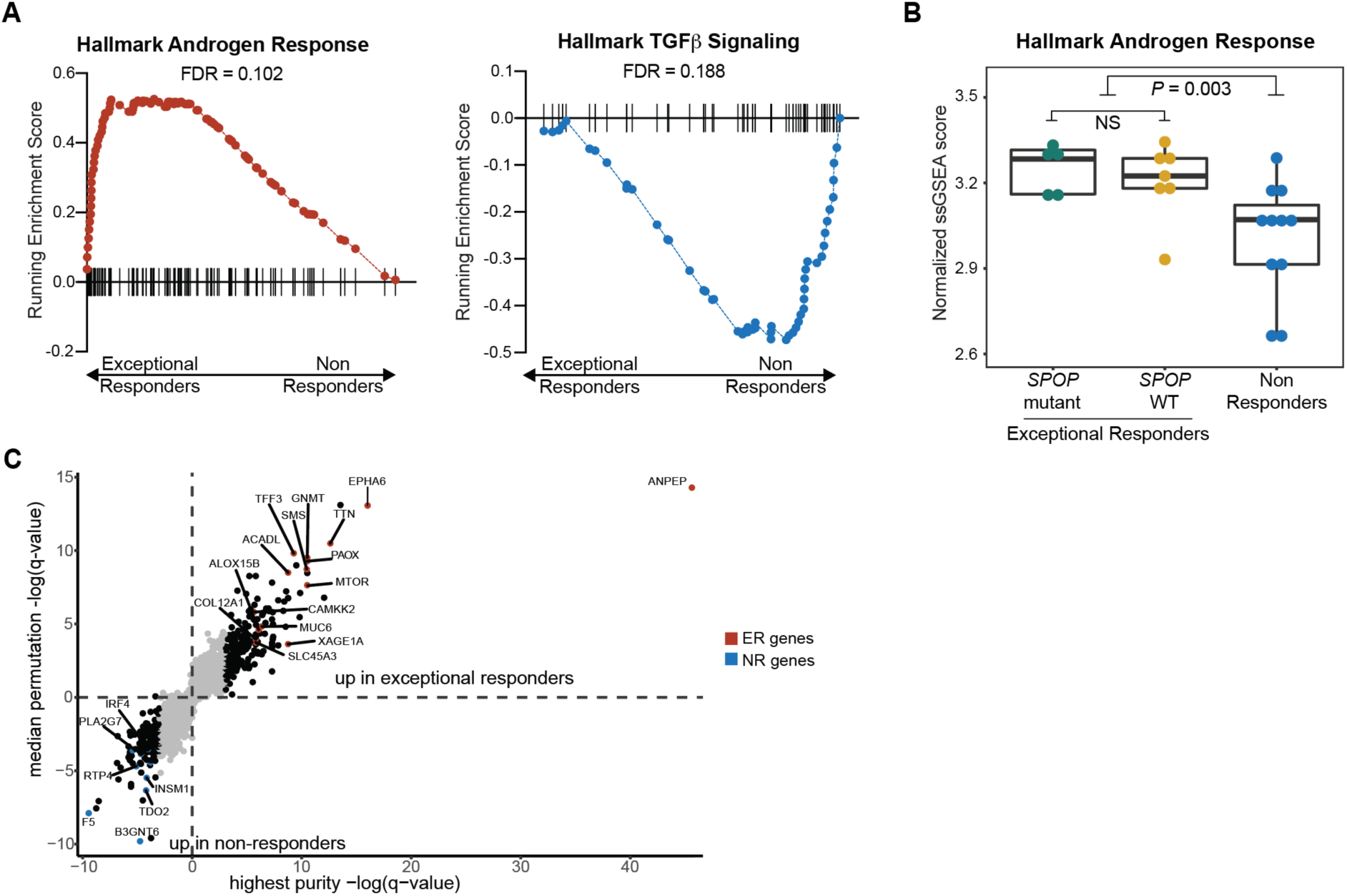
Transcriptional programs of response to neoadjuvant API reveal up-regulated androgen signaling in responders and TGFβ signaling in non-responders. **(A)** GSEA was performed on the highest purity ER versus the highest purity NR samples (Methods). Enrichment plot of hallmark androgen response gene set (FDR = 0.102) and the hallmark TGFβ signaling pathway (FDR = 0.188) are shown here. **(B)** ssGSEA of the hallmark androgen signaling gene set applied to the RNA-seq sample when available from the highest purity sample from each patient, stratified by *SPOP* mutation status (N=5, *SPOP* mutant/ER, N=7, *SPOP* wild type/ER, N=11, NR) and response status. A statistically significant difference was detected in androgen signaling between ER and NR (*P* = 0.003 for ssGSEA score, Wilcoxon-rank sum), but no difference was observed by *SPOP* mutation status amongst ER samples (*P* = 0.935, Wilcoxon-rank sum). **(C)** Differential expression analysis of ER vs NR samples. The y-axis shows the median permutation q-value across random iterations selecting one RNA-seq sample per patient and running edgeR as related to the q-value from edgeR run on RNA-seq data from the highest purity sample from each patient. Genes are colored by response status if they were also differentially expressed by TPM value between ER and NR (*P* < 0.05, Mann-Whitney U test).

## Discussion

In summary, we identified *SPOP* mutation as a truncal event in prostate tumors that have exceptional response to neoadjuvant API in HRLPC, which may have major implications for treatment selection in this high-risk patient population. Exceptional response was also associated with *SPOPL* and *CHD1* copy number loss and additional genomic and transcriptional features, suggesting multiple integrated molecular processes can contribute to exceptional response^27^. *SPOP* is an E3 ligase adaptor protein that forms large oligomers or heteromers with *SPOPL* to promote ubiquitination and proteosomal degradation of target proteins^58^. Mutations in *SPOP* have been proposed to disrupt oligomerization in a dominant negative fashion to reduce substrate ubiquitination, and the loss of *SPOPL* likely leads to a similar outcome^59^. *SPOP* mutation has been associated with longer time on treatment with abiraterone in metastatic castration-resistant prostate cancer^30^, as well as progression-free survival for standard ADT alone in metastatic castration-sensitive prostate cancer^60^. Notably, the genomic profiles of pre-treatment HRLPC samples from an independent neoadjuvant API clinical trial cohort revealed the presence of *SPOP* mutations exclusively in exceptional responders^61^. Our findings, paired with this external validation, demonstrate the translational relevance of mutations in this gene with API in HRLPC.

Consistent with a prior study in preclinical mouse models that identified increased AR and AR-associated transcription factors as a result of *SPOP* mutation^62^, we also identified up-regulated androgen signaling in ER relative to NR as a possible explanation for preferential sensitivity to intensive neoadjuvant API. Our findings are further corroborated mechanistically by (1) Grbesa and colleagues, who showed that *SPOP* mutation is sufficient to reprogram the *AR* towards its oncogenic program in mouse organoids, rendering them more dependent on androgen signaling, and (2) Bernasocchi et al^63^, who described that the expression of *SPOP* mutants in a cell line model of advanced prostate cancer induces greater sensitivity to androgen-deprivation and enzalutamide therapy. Increased androgen signaling was not exclusive to *SPOP* mutated samples. For example, 3 ER patients lacking a *SPOP* mutation were found to have missense mutations in *FOXA1* located at the C-terminal end of the FKHD domain. Mutations in this location were recently characterized^64^ as augmenting the impact of *FOXA1* on *AR* binding to DNA and transcription of target genes, which may explain increased androgen signaling in these non-*SPOP* mutated samples.

Our findings suggest additional prospective molecular stratification may broadly improve selection of neoadjuvant treatment strategies in HRLPC. For example, *PTEN* copy number loss was only observed in NR, consistent with reduced PTEN expression by immunohistochemistry in tumors persisting after treatment on our clinical trial^27^. It is possible that HRLPC patients with *PTEN* loss should not receive neoadjuvant API, but rather may benefit from a different therapy such as *AKT* inhibition as is currently being evaluated with ipatasertib in the IPATential150 trial (NCT03072238)^65^. Furthermore, 2 NR patients had copy number evidence of HRD in the absence of *BRCA* mutation, which may indicate sensitivity to PARP inhibition^52,66^. We also identified enrichment of TGFβ signaling in NR, which parallels the increased TGFβ pathway activity in enzalutamide resistant advanced PCa^67^ and preclinical findings that the TGFβ inhibitor galunisertib augments the anti-tumor activity of enzalutamide^68^. Galunisertib is currently being evaluated in a clinical trial in combination with enzalutamide in advanced PCa (NCT02452008); a similar strategy may be applicable to HRLPC.

The presence of the same *SPOP* mutation within multiple tumor foci from ER patients is in contrast to prior genomic studies of multifocal PCa prostatectomy samples unstratified by treatment^36–40^, which detected *SPOP* mutation either clonally or subclonally in only a subset of sampled regions. It is possible that *SPOP* mutations were present in unsampled foci from NR tumors that respond to therapy, whereas residual tumor foci contain other drivers of resistance to API^69,70^. Integration of multiple molecular features from multifocal biopsies may therefore be required for more accurate treatment selection.

Finally, the small sample size of this cohort and use of whole exome sequencing limits the biomarker generalizability of our findings beyond initial biological discovery. Expanded prospective analyses that include expanded sequencing modalities (e.g. whole genome and transcriptome sequencing) and longer-term follow-up are required to validate *SPOP* mutation, TGFβ pathway activity, alterations in DNA damage repair processes, and androgen signaling as predictive biomarkers for treatment stratification to neoadjuvant API or other approaches in HRLPC. Indeed, studies of these factors and other molecular associations of response are planned as correlative analyses within the international randomized phase 3 PROTEUS study (NCT03767244) of neoadjuvant apalutamide plus ADT in HRLPC. Overall, these findings may ultimately direct molecular patient stratification for the HRLPC patient population most likely to respond to intensive androgen blockade and guide alternative neoadjuvant therapeutic strategies for other molecularly defined subsets of patients.

## METHODS

### Patient Cohort

Patients with localized prostate cancer who were candidates for prostatectomy and met eligibility criteria were offered enrollment on Dana-Farber Cancer Institute IRB approved clinical trial protocols 14-283 (enzalutamide, lupron, abiraterone acetate, and prednisone) or 16-223 (apalutamide, abiraterone acetate, lupron). Patient written and informed consent was obtained for molecular analysis of pre-treatment prostate biopsies under Dana-Farber Cancer Institute IRB approved protocols (01-045/11-104/17-000). All patients who consented for molecular analysis and met criteria for exceptional response (either complete response - no tumor tissue upon pathologic review at time of prostatectomy, or minimal residual disease - < 5 mm of tumor tissue in longest dimension at time of prostatectomy) or non-response (pathologic T3 or lymph node positivity at time of prostatectomy) were included. A total of 32 patients consented, had available tissue, and had nucleic acid successfully extracted from tumor tissue. Eight patients were ultimately excluded from final analysis because of initial incorrect response classification (1 patient – actually an intermediate responder), or sequencing/downstream QC issues (7 patients – metrics detailed below), resulting in a final cohort of 24 patients.

### Whole exome sequencing and whole transcriptome sequencing

DNA extraction, library preparation and WES were performed for samples as previously described^47^. Each WES sample consisted of DNA extracted from the tumor tissue in one prostate biopsy core. Slides were cut from FFPE biopsy blocks, macrodissected for tumor-enriched tissue, and deparaffinized. DNA and RNA extraction were performed using the QIAGEN AllPrep FFPE DNA/RNA Extraction kit. Germline DNA was obtained from peripheral blood mononuclear cells. WES libraries were generated from 100 ng of starting DNA material. After processing and size selection, samples were subjected to exonic hybrid capture, then sequenced using the Illumina HiSeq platform.

Total RNA was assessed for quality and the percentage of fragments with a size greater than 200 nucleotides (DV200) was calculated using software. cDNA library synthesis and capture were performed using the Illumina TruSeq RNA Access Library Prep kit (now known as Illumina TruSeq RNA Exome kit). Amplified libraries were quantified using an automated PicoGreen assay. Flowcell cluster amplification and sequencing were performed according to the manufacturer’s protocols using the Illumina NextSeq 500 platform. Each run generated 76 bp stranded paired end reads. Raw sequencing data were processed using the Broad Picard Pipeline, which includes demultiplexing and data aggregation.

### WES Quality Control

Samples were included for analysis only if they successfully underwent WES, met criteria for either exceptional or non-response, had a matched germline normal sample, and met our joint quality control criterion. To estimate cross sample contamination we used ContEst^71^, and applied a cutoff of <= 4%. The median cross sample contamination was 0.1%. We utilized the GATK3.7 DepthOfCoverage tool^72^ to ascertain the mean target coverage for tumor and normal samples, and required at least 50x coverage for the tumor sample and 30x coverage for the corresponding normal sample. Finally, we utilized both FACETS^73^ and ABSOLUTE^74^ to determine tumor purity and excluded samples with tumor purity < 20% by both algorithms.

### Variant Calling and Mutational Significance Analysis

Reads were aligned using BWA^75^ v0.5.9 and somatic mutations called using a customized version of the Getz Lab CGA WES Characterization pipeline (https://portal.firecloud.org/#methods/getzlab/CGA_WES_Characterization_Pipeline_v0.1_Dec2 018/) developed at the Broad Institute. We used ContEst^71^ to estimate cross sample contamination. We used MuTect^76^ v1.1.6 to call single nucleotide variants, and Strelka^77^ v1.0.11 to call indels. MuTect2.1^78^ was used to confirm Strelka indel calls. MuTect v1.1.6 output was filtered for FFPE^72^ and 8-oxoG^79^ sequencing artifacts using GATK FilterByOrientationBias. DeTiN^80^ was used to rescue true somatic variants that were removed due to tumor-in-normal contamination. Variant calls were subsequently filtered through a panel of normal samples to remove artifacts from miscalled germline alterations and other rare error modes. Variants were annotated using VEP, Oncotator, and vcf2maf v1.6.17 (https://github.com/mskcc/vcf2maf). The variant allele frequency of detected mutations that passed QC steps is indicated in Table S2.

To identify significantly mutated genes across our cohort, we utilized MutSigCV2^34^, utilizing the highest purity sample per patient since WES data from intra-tumoral foci are not independent of each other.

### Copy number calling and significance

Allelic copy number, tumor purity and tumor ploidy were analyzed using both FACETS^73^ and ABSOLUTE^74^. Purity and ploidy calls by each algorithm were generally concordant (spearman ρ = 0.707, *P* = 3.9×10^-8^, Supplemental Table 2). Copy number alterations (CNA), purity, ploidy, and whole genome doubling status used for analyses in Figure 1 were based on FACETS calls, since FACETS provides allelic copy number calls for the X chromosome. We utilized GISTIC 2.0^33^ to identify focal regions with significant enrichment of amplifications and deletions using genome segmentation files generated by GATK 3.7. Again, we restricted the analysis to the highest purity sample from each patient. To remove germline noise, we ran GISTIC2.0 on the merged matched normal segmentation file and removed significantly enriched regions using amplification and deletion thresholds of 0.1. Any germline region with a q-value < 0.25 (default) was considered significant and excluded from the somatic analysis. To identify focal regions with significant enrichment of somatic amplifications/deletions, we utilized amplification and deletion thresholds of 0.3, and applied a q-value cutoff of < 0.1.

### Phylogenetic analysis

A union set of mutations across all samples from each patient was generated and force calling was performed to assess the variant allele fraction of each mutation within each sample. The cancer cell fraction (CCF) of mutations were defined using ABSOLUTE^74^, which calculates the CCF based on allelic fraction, purity, and local copy number. To reconstruct the clonal architecture of prostate cancer tumors, we used the PhylogicNDT^41^ Cluster module, which was initialized using ABSOLUTE output and utilizes Dirichlet clustering to determine the number of clusters and the respective assignment of each mutation to a cell subpopulation (or subclone). The CCF annotated MAF file from ABSOLUTE and tumor purity for each WES sample per patient were used as inputs to the clustering method. The outputs from the PhylogicNDT Cluster were then used as inputs to the PhylogicNDT BuildTree module, which produces a series of phylogenetic trees ordered by likelihood, with the baseline assumption that all biopsies included in the analysis are related. The phylogenetic trees with the highest likelihood were used in the analyses of this study. The cluster average CCFs and individual mutation CCFs are shown in Table S7.

### Mutational signature and homologous recombination deficiency analysis

Mutational processes in our cohort were determined using deconstructSigs^46^ using default parameters with COSMIC^50^ v2 signatures as the reference. The highest purity sample per patient was used for this analysis. A signature was assessed as present if the signature contribution was greater than 6%. Samples were annotated for the presence or absence of each signature, and enrichment of each signature by response was determined by Fisher’s exact test. We also utilized SigMA^47^, which leverages multiple methods including likelihood-based statistics to classify known signatures across cancer types. For this analysis, we utilized default parameters with the exception of setting the tumor type to “prost”, the data parameter set to “seqcap”, and the check_msi parameter was set to “true”. Signature 3 status was determined using the category classification output by SigMa; however, we also utilized the signature 3 likelihood output to identify high-confidence signature 3 tumors.

To calculate the number of loss of heterozygosity, telomeric allelic imbalance and large scale transition events, we used FACETS allelic copy number calls for the highest purity sample per patient as input into the scarHRD^48^ package (https://github.com/sztup/scarHRD). Differences in scarHRD score by response were assessed by the Wilcoxon rank-sum test.

### Germline variant discovery

To call short germline single-nucleotide polymorphisms, insertions, and deletions from germline WES data, we used DeepVariant^81^ (v0.8.0). Specifically, we used the publicly released WES model (https://console.cloud.google.com/storage/browser/deepvariant/models/DeepVariant/0.8.0/Deep Variant-inception_v3-0.8.0+data-wes_standard/) to generate single-sample germline variant call files using the human genome reference GRCh37(b37). We filtered variants with bcftools v1.9 to only keep high-quality variants annotated as “PASS” in the “FILTER” column. The high-quality variants were merged into single-sample Variant Call Format (VCF) files using the “ “CombineVariants” module of GATK 3.7 (https://github.com/broadinstitute/gatk/releases). To decompose multiallelic variants and normalize variants, we used the computational package vt v3.13 (https://github.com/atks/vt). Lastly, germline variants were annotated using the VEP v92 with the publicly released GRCh37 cache file (https://github.com/Ensembl/ensembl-vep).

### RNA-seq processing and normalization

After sequencing, adapters were trimmed with cutadapt^82^ v2.2 and reads were aligned using STAR^83^ aligner v2.7.2b. We used multi-sample 2-pass mapping for all samples from each patient, first mapping all samples, merging the SJ.out.tab files, then running the second pass with the jointly called splice junctions. STAR-aligned bams were passed into RSEM^84^ to generate gene-level transcript counts and transcript per million (TPM) quantifications using the Gencode30 reference. STAR chimeric junctions were supplied to STAR-Fusion^85^ v1.7.0 in 562 kickstart mode to call *TMPRSS2*-*ERG* fusions.

For RNA-seq quality control, sequencing- and alignment-specific metrics were examined for each sample. The following alignment metrics were considered and examined for outliers: number and percentage of uniquely mapped reads, number of high-quality reads, intronic rate, intergenic rate and rRNA rate. Samples were clustered across quality-control metrics using principal-component analysis that incorporated rates of high-quality reads in exonic, intronic, intragenic and intragenic regions as well as base mismatch rate, unique mapping rate, and rRNA rate. Two samples were excluded based on consistently being outliers across various metrics as well as by principal component analysis. Finally, only transcriptomes from tumors whose WES also passed quality control were included. One sample with WES data did not have an RNA-seq sample that successfully completed sequencing, resulting in a total of 43 RNA-seq samples in our analysis cohort.

### Gene set enrichment analysis

GSEA^53^ was performed using the Cancer Hallmarks gene sets from MSigDB (https://cloud.genepattern.org/). The trimmed mean of m-values (TMM) normalized counts for RefSeq genes as generated by edgeR^86^ were used as inputs. Genes with less than 5 reads across all samples were excluded from analysis. We used default settings with 1,000 phenotype permutations to generate *P* and *Q* values and compared the highest purity RNA-seq samples between ER and NR. A gene set was considered significant with a false discovery rate of less than 25%. To generate nonparametric gene set scores for the hallmark androgen signaling gene set, we generated ssGSEA projections^55^ using rank normalization. Differences in ssGSEA score were assessed by the Wilcoxon rank-sum test.

### Differential expression analysis

RNA differential expression analysis was completed using edgeR^86,87^ v3.11 with the gene level counts from RSEM as input and tumor purity as a covariate in the design matrix. We performed differential expression analysis between ER and NR patients using two approaches. We first performed differential expression using the highest purity sample from each patient. To determine the generalizability of these findings across the cohort and leverage the sequencing of multiple samples, we also undertook a sampling approach. We randomly selected one tumor sample from each patient to include in an iteration of differential expression analysis; we repeated this process 1000 times and recorded the median edgeR *Q*-value of each gene across the 1000 iterations. To ensure that differential expression results were not driven by outliers, we also conducted a one-sided Mann-Whitney U test on the gene level TPMs. Genes were considered differentially expressed if they had a *Q* value below 0.05 in the highest purity analysis, a *Q* value below 0.05 in the sampling approach, and a *P* value below 0.05 in the Mann-Whitney U test.

## STATISTICAL ANALYSIS

The statistical details of all analyses are reported in the main text, figure legends, and figures, which includes the statistical test performed and statistical significance.

## DATA AVAILABILITY

Raw sequencing data will be available at dbGAP accession phs001988.v1.p1at the time of publication.

## ACKNOWLEDGEMENTS

This work was supported by NIH T32 CA009172 (A.K.T.), NCI F31CA239347 (J.R.C.), PCF Young Investigator Award YIA18YOUN02 (S.H.A), American Society of Clinical Oncology (ASCO) Conquer Cancer Foundation Career Development Award CDA13167 (S.H.A.), NIH T32 GM008313 and NSF GRFP DGE1144152 (M.X.H.), PCF-V Foundation Challenge Award (E.M.V.), PCF-Movember Challenge Award (E.M.V., M.E.T.), NIH U01 CA233100 (E.M.V.), NIH R37 CA222574 (E.M.V.), NIH R01 CA227388 (E.M.V.), Mark Foundation Emerging Leader Award (E.M.V.), and clinical trial support from Pfizer and Janssen. We are incredibly grateful for the patients who made this study possible.

## AUTHOR CONTRIBUTIONS

M.-E.T., and E.M.V.A designed the overall study. Z.Z., R.M, M-E.T. and E.M.V.A. supervised and organized the sample collection and preparation. A.K.T., A.T.M.C., J.C., J.R.C., S.A.W., J.P. A.B.M., and M.X.H. contributed to the analysis of WES and RNA-seq data. A.K.T., J.R.C., J.P., D.L., I.L, and G.G. specifically contributed to phylogenetic analysis of samples. A.K.T., S.C. and S.H.A. analyzed germline sequencing data. A.K.T., A.T.M.C., J.C. J.R.C., M-E.T. and E.M.V.A. contributed to interpretation of results and manuscript preparation. All authors reviewed and approved the final manuscript.

---

**Supplementary Figure 1:**
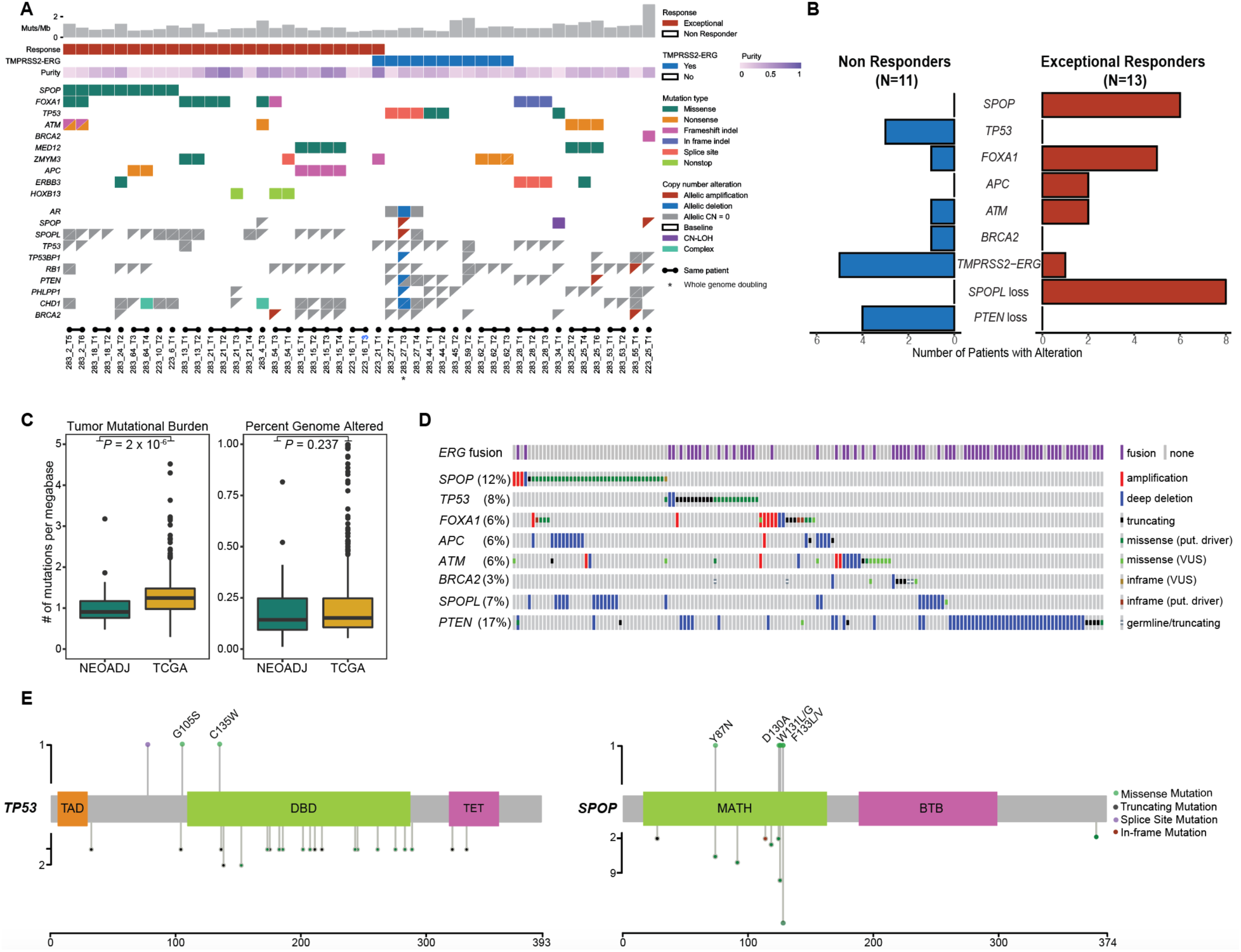
Genomic characteristics of all included exomes and comparison to TCGA samples. **(A)** Co-mutation plot for all included exomes illustrating the patient, select single nucleotide and insertion/deletion events, as well as CNA and *TMPRSS2-ERG* RNA fusion status. Each row represents the mutation or copy number status for the indicated gene, and each column represents a patient sample. **(B)** Count of selected molecular alterations present in highest purity sample per patient. All alterations were present in each sample assayed per patient with the exception of *SPOPL* copy number loss in patient 283_21. **(C)** Comparison of TMB (*P* = 2 x 10^-6^, Wilcoxon rank-sum) and PGA (*P* = 0.237, Kolmogorov-Smirnov) between this cohort (n = 24) and treatment unselected patients from the TCGA (n=333) patients. **(D)** cBioPortal^89,90^ OncoPrint of molecular alterations from Supplemental Figure 1b within TCGA localized prostate cancer samples. **(e)** Lollipop plots of detected alterations in *TP53* and *SPOP* to specific genomic changes in neoadjuvantly treated patients (top) versus TCGA changes (bottom). Lollipop height is proportional to number of alterations detected.

**Supplementary Figure 2:**
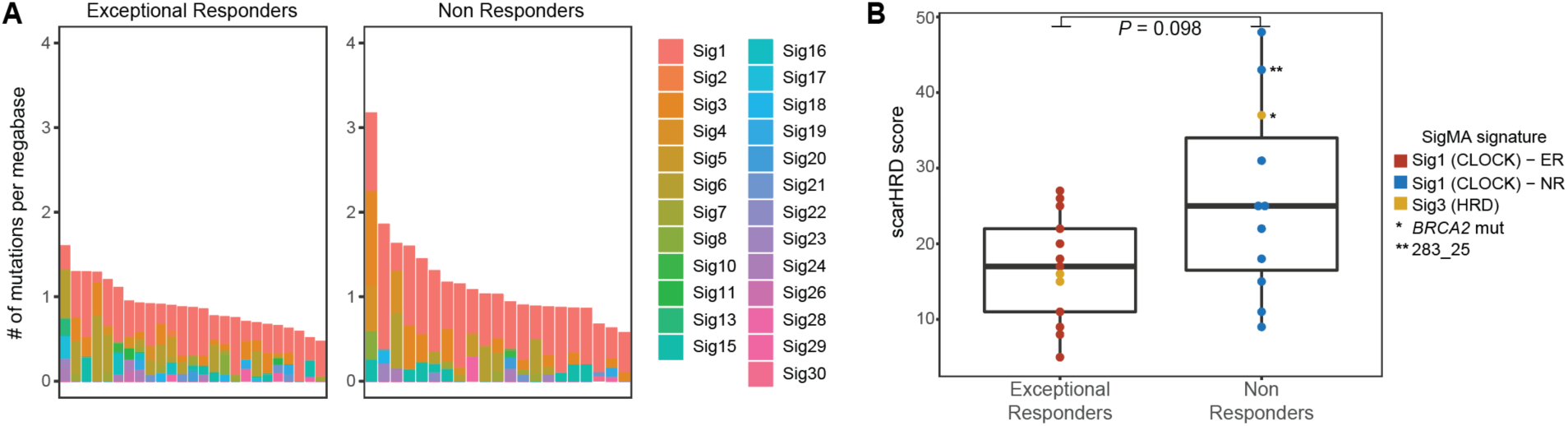
Mutational processes present in neoadjuvantly treated samples stratified by response. **(A)** deConstructSigs output for highest purity ER and NR samples per patient. The height of each bar is colored by the proportion of mutations detected attributed to each of the COSMIC v2 mutational signatures. Differences in mutational signatures were assessed between ER and NR by Fisher’s exact test. **(B)** scarHRD output for copy-number based evidence of HRD between highest purity ER (N = 13) and NR (N = 11). Samples with evidence of signature-3 mutational process by SigMA are colored in yellow, the rest of the samples are colored by response and were identified as predominantly having the signature-1 mutational process. Difference in scarHRD score (*P* = 0.098) was assessed by Wilcoxon-rank sum.

**Supplemental Figure 3:**
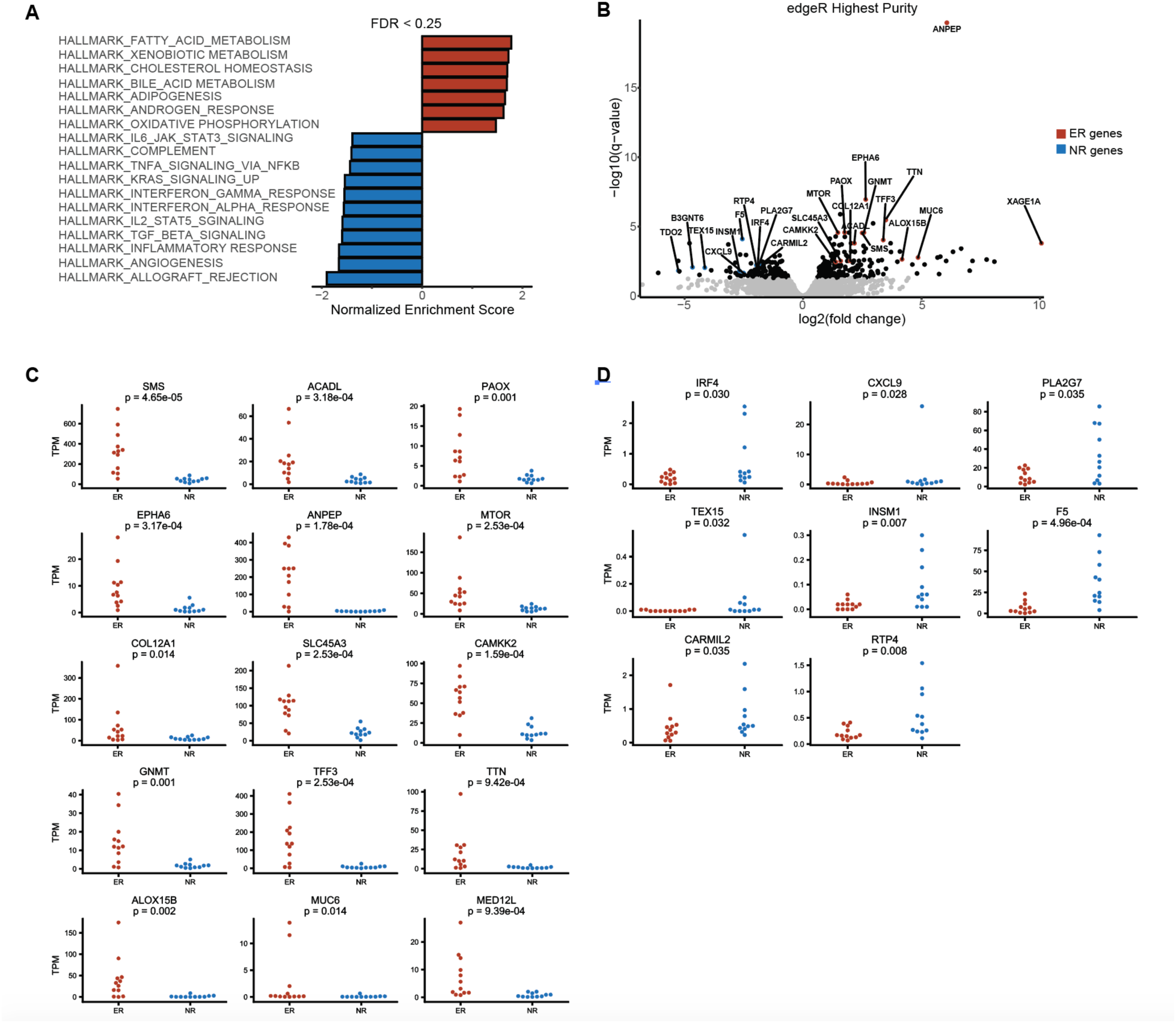
Gene sets and individual genes enriched in ER versus NR patients. **(A)** Gene set enrichment results for all Hallmark MSigDB gene sets enriched in each response category (ER = red, NR = blue) with a false discovery rate of less than 25%. **(B)** edgeR plot of fold change versus significance for RNA-seq samples from the highest purity biopsy per each patient. Genes significantly enriched in ER by both edgeR q-value < 0.05 and by difference in TPM (Mann-Whitney U test < 0.05) are colored red. Genes significantly enriched in NR by both edgeR q-value < 0.05 and by difference in TPM (Mann-Whitney U test < 0.05) are colored blue **(C)** TPM from highest purity sample for RNA-seq expression data for selected genes significantly enriched in ER samples by both TPM and edgeR comparison. **(D)** TPM from highest purity sample for RNA-seq expression data for selected genes significantly enriched in NR samples by both TPM and edgeR comparison.

## Supplemental Tables

**Table S1:** Final cohort clinical and genomic characteristics

**Table S2:** Union set of forcecalled nonsynonymous mutations

**Table S3:** GISTIC output

**Table S4:** Significance testing for detected mutations

**Table S5:** Mutational processes as assessed by deConstructSigs, scarHRD, and SigMA

**Table S6:** Frequency of selected germline alterations by response

**Table S7:** PhylogicNDT mutational clustering output

**Table S8:** RNA-seq RSEM generated counts for RefSeq genes

## REFERENCES

1. Eggener, S. E. et al. Predicting 15-year prostate cancer specific mortality after radical prostatectomy. J. Urol. 185, 869–875 (2011).

2. Siegel, R. L., Miller, K. D. & Jemal, A. Cancer statistics, 2020. *CA*. Cancer J. Clin. 70, 7–30 (2020).

3. Kelly, S. P., Anderson, W. F., Rosenberg, P. S. & Cook, M. B. Past, Current, and Future Incidence Rates and Burden of Metastatic Prostate Cancer in the United States. Eur. Urol. Focus 4, 121–127 (2018).

4. Cooperberg, M. R., Broering, J. M. & Carroll, P. R. Time Trends and Local Variation in Primary Treatment of Localized Prostate Cancer. J. Clin. Oncol. 28, 1117–1123 (2010).

5. Stephenson, A. J. et al. Prostate cancer-specific mortality after radical prostatectomy for patients treated in the prostate-specific antigen era. J. Clin. Oncol. 27, 4300–4305 (2009).

6. Grossman, H. B. et al. Neoadjuvant Chemotherapy plus Cystectomy Compared with Cystectomy Alone for Locally Advanced Bladder Cancer. N. Engl. J. Med. 349, 859–866 (2003).

7. Petrelli, F. et al. Correlation of pathologic complete response with survival after neoadjuvant chemotherapy in bladder cancer treated with cystectomy: A meta-analysis. Eur. Urol. 65, 350–357 (2014).

8. Cortazar, P. et al. Pathological complete response and long-term clinical benefit in breast cancer: The CTNeoBC pooled analysis. Lancet 384, 164–172 (2014).

9. Symmans, W. F. et al. Measurement of residual breast cancer burden to predict survival after neoadjuvant chemotherapy. J. Clin. Oncol. 25, 4414–4422 (2007).

10. Gianni, L. et al. 5-year analysis of neoadjuvant pertuzumab and trastuzumab in patients with locally advanced, inflammatory, or early-stage HER2-positive breast cancer (NeoSphere): a multicentre, open-label, phase 2 randomised trial. Lancet Oncol. 17, 791–800 (2016).

11. Fayanju, O. M. et al. The Clinical Significance of Breast-only and Node-only Pathologic Complete Response (pCR) After Neoadjuvant Chemotherapy (NACT): A Review of 20,000 Breast Cancer Patients in the National Cancer Data Base (NCDB). Ann. Surg. 268, (2018).

12. Ryan, S. T., Patel, D. N., Parsons, J. K. & McKay, R. R. Neoadjuvant Approaches Prior to Radical Prostatectomy. Cancer J. (United States) 26, 2–12 (2020).

13. McKay, R. R., Choueiri, T. K. & Taplin, M. E. Rationale for and review of neoadjuvant therapy prior to radical prostatectomy for patients with high-risk prostate cancer. Drugs 73, 1417–1430 (2013).

14. L., T. et al. Systematic Review of Systemic Therapies and Therapeutic Combinations with Local Treatments for High-risk Localized Prostate Cancer. Eur. Urol. 75, 44–60 (2019).

15. Mostaghel, E. A. et al. Intraprostatic androgens and androgen-regulated gene expression persist after testosterone suppression: Therapeutic implications for castration-resistant prostate cancer. Cancer Res. 67, 5033–5041 (2007).

16. Febbo, P. G. et al. Genomic strategy for targeting therapy in castration-resistant prostate cancer. J. Clin. Oncol. 27, 2022–2029 (2009).

17. Taplin, M.-E. et al. Intense Androgen-Deprivation Therapy With Abiraterone Acetate Plus Leuprolide Acetate in Patients With Localized High-Risk Prostate Cancer: Results of a Randomized Phase II Neoadjuvant Study. J. Clin. Oncol. 32, 3705–3715 (2014).

18. De Bono, J. S. et al. Abiraterone and Increased Survival in Metastatic Prostate Cancer. N. Engl. J. Med. 364, 1995–2005 (2011).

19. Ryan, C. J. et al. Abiraterone in metastatic prostate cancer without previous chemotherapy. N. Engl. J. Med. 368, 138–48 (2013).

20. Fizazi, K. et al. Abiraterone plus Prednisone in Metastatic, Castration-Sensitive Prostate Cancer. N. Engl. J. Med. 377, 352–60 (2017).

21. James, N. D. et al. Abiraterone for Prostate Cancer Not Previously Treated with Hormone Therapy. N. Engl. J. Med. 377, 338–351 (2017).

22. Scher, H. I. et al. Increased survival with enzalutamide in prostate cancer after chemotherapy. N. Engl. J. Med. 367, 1187–97 (2012).

23. Hussain, M. et al. Enzalutamide in Men with Nonmetastatic, Castration-Resistant Prostate Cancer. N. Engl. J. Med. 278, 2465–74 (2018).

24. Davis, I. D. et al. Enzalutamide with Standard First-Line Therapy in Metastatic Prostate Cancer. N. Engl. J. Med. NEJMoa1903835 (2019). doi:10.1056/NEJMoa1903835

25. Smith, M. R. et al. Apalutamide treatment and metastasis-free survival in prostate cancer. N. Engl. J. Med. 378, 1408–1418 (2018).

26. Chi, K. N. et al. Apalutamide for metastatic, castration-sensitive prostate cancer. N. Engl. J. Med. 381, 13–24 (2019).

27. McKay, R. R. et al. Evaluation of intense androgen deprivation before prostatectomy: A randomized Phase II trial of enzalutamide and leuprolide with or without abiraterone. J. Clin. Oncol. 37, 923–931 (2019).

28. McKay, R. R. et al. Post prostatectomy outcomes of patients with high-risk prostate cancer treated with neoadjuvant androgen blockade. Prostate Cancer Prostatic Dis. 21, 364–372 (2018).

29. Abida, W. et al. Genomic correlates of clinical outcome in advanced prostate cancer. Proc. Natl. Acad. Sci. 201902651 (2019). doi:10.1073/pnas.1902651116

30. Boysen, G. et al. SpoP-mutated/CHD1-deleted lethal prostate cancer and abiraterone sensitivity. Clin. Cancer Res. 24, 5585–5593 (2018).

31. Klein, E. A. et al. A genomic classifier improves prediction of metastatic disease within 5 years after surgery in node-negative high-risk prostate cancer patients managed by radical prostatectomy without adjuvant therapy. Eur. Urol. 67, 778–786 (2015).

32. Eggener, S. E. et al. Molecular biomarkers in localized prostate cancer: ASCO guideline. J. Clin. Oncol. 38, 1474–1494 (2020).

33. Mermel, C. H. et al. GISTIC2.0 facilitates sensitive and confident localization of the targets of focal somatic copy-number alteration in human cancers. Genome Biol. 12, 1–14 (2011).

34. Lawrence, M. S. et al. Mutational heterogeneity in cancer and the search for new cancer-associated genes. Nature 499, 214–218 (2013).

35. Abeshouse, A. et al. The Molecular Taxonomy of Primary Prostate Cancer. Cell 163, 1011–1025 (2015).

36. Løvf, M. et al. Multifocal Primary Prostate Cancer Exhibits High Degree of Genomic Heterogeneity. Eur. Urol. 75, 498–505 (2019).

37. Wei, L. et al. Intratumoral and Intertumoral Genomic Heterogeneity of Multifocal Localized Prostate Cancer Impacts Molecular Classifications and Genomic Prognosticators. Eur. Urol. 71, 183–192 (2017).

38. Boutros, P. C. et al. Spatial genomic heterogeneity within localized, multifocal prostate cancer. Nat. Genet. 47, 736–745 (2015).

39. Fraser, M. et al. Genomic hallmarks of localized, non-indolent prostate cancer. Nature 541, 359–364 (2017).

40. Espiritu, S. M. G. et al. The Evolutionary Landscape of Localized Prostate Cancers Drives Clinical Aggression. Cell 173, (2018).

41. Leshchiner, I. et al. Comprehensive analysis of tumour initiation, spatial and temporal progression under multiple lines of treatment. bioRxiv (2018). doi:https://doi.org/10.1101/508127

42. Gruber, M. et al. Growth dynamics in naturally progressing chronic lymphocytic leukaemia. Nature (2019). doi:10.1038/s41586-019-1252-x

43. Boysen, G. et al. SPOP mutation leads to genomic instability in prostate cancer. Elife 4, 1–18 (2015).

44. Hjorth-Jensen, K. et al. SPOP promotes transcriptional expression of DNA repair and replication factors to prevent replication stress and genomic instability. Nucleic Acids Res. 46, 9484–9495 (2018).

45. Niu, B. et al. MSIsensor: Microsatellite instability detection using paired tumor-normal sequence data. Bioinformatics 30, 1015–1016 (2014).

46. Rosenthal, R., McGranahan, N., Herrero, J., Taylor, B. S. & Swanton, C. deconstructSigs: Delineating mutational processes in single tumors distinguishes DNA repair deficiencies and patterns of carcinoma evolution. Genome Biol. 17, 1–11 (2016).

47. Gulhan, D. C., Lee, J. J. K., Melloni, G. E. M., Cortés-Ciriano, I. & Park, P. J. Detecting the mutational signature of homologous recombination deficiency in clinical samples. Nat. Genet. 51, 912–919 (2019).

48. Sztupinszki, Z. et al. Migrating the SNP array-based homologous recombination deficiency measures to next generation sequencing data of breast cancer. npj Breast Cancer 4, 8–11 (2018).

49. Alexandrov, L. B. et al. The repertoire of mutational signatures in human cancer. Nature 578, 94–101 (2020).

50. Tate, J. G. et al. COSMIC: The Catalogue Of Somatic Mutations In Cancer. Nucleic Acids Res. 47, D941–D947 (2019).

51. Telli, M. L. et al. Homologous Recombination Deficiency (HRD) Score Predicts Response to Platinum-Containing Neoadjuvant Chemotherapy in Patients with Triple-Negative Breast Cancer. Clin. Cancer Res. 22, 3764–3773 (2016).

52. Sztupinszki, Z. et al. Detection of Molecular Signatures of Homologous Recombination Deficiency in Prostate Cancer with or without BRCA1/2 Mutations. Clin. Cancer Res. 2673–2681 (2020). doi:10.1158/1078-0432.ccr-19-2135

53. Subramanian, A. et al. Gene set enrichment analysis: a knowledge-based approach for interpreting genome-wide expression profiles. Proc. Natl. Acad. Sci. U. S. A. 102, 15545–50 (2005).

54. Liberzon, A. et al. The Molecular Signatures Database (MSigDB) hallmark gene set collection. Cell Syst. 1, 417–425 (2015).

55. Barbie, D. A. et al. Systematic RNA interference reveals that oncogenic KRAS-driven cancers require TBK1. Nature 462, 108–112 (2009).

56. Sørensen, K. D. et al. Prognostic significance of aberrantly silenced ANPEP expression in prostate cancer. Br. J. Cancer 108, 420–428 (2013).

57. Vainio, P. et al. Phospholipase PLA2G7, associated with aggressive prostate cancer, promotes prostate cancer cell migration and invasion and is inhibited by statins. Oncotarget 2, 1176–1190 (2011).

58. Errington, W. J. et al. Adaptor protein self-assembly drives the control of a cullin-RING ubiquitin ligase. Structure 20, 1141–1153 (2012).

59. Cuneo, M. J. & Mittag, T. The ubiquitin ligase adaptor SPOP in cancer. FEBS J. 286, 3946–3958 (2019).

60. Swami, U. et al. Association of SPOP Mutations with Outcomes in Men with De Novo Metastatic Castration-sensitive Prostate Cancer. Eur. Urol. (2020). doi:10.1016/j.eururo.2020.06.033

61. Wilkinson, S. et al. Nascent Prostate Cancer Heterogeneity Drives Evolution and Resistance to Intense Hormonal Therapy. Eur. Urol. 1–12 (2021). doi10.1016/j.eururo.2021.03.009

62. Blattner, M. et al. SPOP Mutation Drives Prostate Tumorigenesis In Vivo through Coordinate Regulation of PI3K/mTOR and AR Signaling. Cancer Cell 31, 436–451 (2017).

63. Bernasocchi, T. et al. Dual functions of SPOP and ERG dictate androgen therapy responses in prostate cancer. Nat. Commun. 12, 1–18 (2021).

64. Parolia, A. et al. Distinct structural classes of activating FOXA1 alterations in advanced prostate cancer. Nature (2019). doi:10.1038/s41586-019-1347-4

65. J.S. de Bono, S. Bracarda, C.N. Sternberg, K.N. Chi, D. Olmos, S. Sandhu, C. Massard, N. Matsubara, B. Alekseev, R. Gafanov, F. Parnis, G.L. Buchschacher Jr, L. Corrales, M. Borre, G. Vasconcelos Alves, J. Garcia, M. Harle-Yge, G. Chen, M.J. Wongchenko, C. S. LBA4 - IPATential150: Phase III study of ipatasertib (ipat) plus abiraterone (abi) vs placebo (pbo) plus abi in metastatic castration-resistant prostate cancer (mCRPC). Ann. Oncol. 31, S1142–S1215 (2020).

66. De Bono, J. et al. Olaparib for metastatic castration-resistant prostate cancer. N. Engl. J. Med. 382, 2091–2102 (2020).

67. He, M. X. et al. Transcriptional mediators of treatment resistance in lethal prostate cancer. bioRxiv 2020.03.19.998450 (2020). doi:10.1101/2020.03.19.998450

68. Paller, C. et al. TGF-β receptor I inhibitor enhances response to enzalutamide in a pre-clinical model of advanced prostate cancer. Prostate 79, 31–43 (2019).

69. Sowalsky, A. G. et al. Neoadjuvant-intensive androgen deprivation therapy selects for prostate tumor foci with diverse subclonal oncogenic alterations. Cancer Res. 78, 4716–4730 (2018).

70. Wilkinson, S. et al. A case report of multiple primary prostate tumors with differential drug sensitivity. Nat. Commun. 11, 1–8 (2020).

71. Cibulskis, K. et al. ContEst: Estimating cross-contamination of human samples in next-generation sequencing data. Bioinformatics 27, 2601–2602 (2011).

72. Van der Auwera, G. A. et al. From FastQ data to high confidence variant calls: the Genome Analysis Toolkit best practices pipeline. Curr. Protoc. Bioinforma. 43, 11.10.1–11.10.33 (2013).

73. Shen, R. & Seshan, V. E. FACETS: Allele-specific copy number and clonal heterogeneity analysis tool for high-throughput DNA sequencing. Nucleic Acids Res. 44, 1–9 (2016).

74. Carter, S. L. et al. Absolute quantification of somatic DNA alterations in human cancer. Nat. Biotechnol. 30, 413–421 (2012).

75. Li, H. & Durbin, R. Fast and accurate short read alignment with Burrows–Wheeler transform. Bioinformatics 25, 1754–1760 (2009).

76. Cibulskis, K. et al. Sensitive detection of somatic point mutations in impure and heterogeneous cancer samples. Nat. Biotechnol. 31, 213–219 (2013).

77. Saunders, C. T. et al. Strelka: Accurate somatic small-variant calling from sequenced tumor-normal sample pairs. Bioinformatics 28, 1811–1817 (2012).

78. Benjamin, D. et al. Calling Somatic SNVs and Indels with Mutect2. bioRxiv 861054 (2019). doi:10.1101/861054

79. Costello, M. et al. Discovery and characterization of artifactual mutations in deep coverage targeted capture sequencing data due to oxidative DNA damage during sample preparation. Nucleic Acids Res. 41, 1–12 (2013).

80. Taylor-Weiner, A. et al. DeTiN: Overcoming tumor-in-normal contamination. Nat. Methods 15, 531–534 (2018).

81. Poplin, R. et al. A universal SNP and small-indel variant caller using deep neural networks. Nat. Biotechnol. 36, 983–987 (2018).

82. Martin, M. Cutadapt removes adapter sequences from high-throughput sequencing reads. EMBnet.journal*; Vol* 17*, No* *1* Next Gener. Seq. Data Anal. (2011). doi:10.14806/ej.17.1.200

83. Dobin, A. et al. STAR: ultrafast universal RNA-seq aligner. Bioinformatics 29, 15–21 (2013).

84. Nowell, P. C. Linked references are available on JSTOR for this article : The Clonal Evolution of Tumor Cell Populations. Science (80-.). 194, 23–28 (1976).

85. Haas, B. J. et al. Accuracy assessment of fusion transcript detection via read-mapping and de novo fusion transcript assembly-based methods. Genome Biol. 20, 1–16 (2019).

86. McCarthy, D. J., Chen, Y. & Smyth, G. K. Differential expression analysis of multifactor RNA-Seq experiments with respect to biological variation. Nucleic Acids Res. 40, 4288–4297 (2012).

87. Robinson, M. D., McCarthy, D. J. & Smyth, G. K. edgeR: a Bioconductor package for differential expression analysis of digital gene expression data. Bioinformatics 26, 139–140 (2009).

88. Crowdis, J., He, M. X., Reardon, B. & Van Allen, E. M. CoMut: Visualizing integrated molecular information with comutation plots. Bioinformatics (2020). doi:10.1093/bioinformatics/btaa554

89. Cerami, E. et al. The cBio cancer genomics portal: an open platform for exploring multidimensional cancer genomics data. Cancer Discov. 2, 401–4 (2012).

90. Jianjiong, G. et al. Integrative analysis of complex cancer genomics and clinical profiles using the cBioPortal. Sci. Signal. 6, pl1 (2013).

